# Probiotic Supplementation Reduces Inflammatory Profiles But Does Not Prevent Oral Immune Perturbations During SIV Infection

**DOI:** 10.1101/2020.12.17.423321

**Authors:** Rhianna Jones, Kyle Kroll, Courtney Broedlow, Luca Schifanella, Scott Smith, Brady Hueber, Spandan V. Shah, Daniel R. Ram, Cordelia Manickam, Valerie Varner, Nichole R. Klatt, R. Keith Reeves

## Abstract

HIV/SIV infections lead to massive loss of mucosal CD4+ T cells and breakdown of the epithelial mucosa resulting in severe microbial dysbiosis and chronic immune activation that ultimately drive disease progression. Moreover, disruption of one of the most understudied mucosal environments, the oral cavity, during HIV-induced immunosuppression results in significant microbial and neoplastic co-morbidities and contributes to and predicts distal disease complications. In this study we evaluated the effects of oral probiotic supplementation (Pbx), which can stimulate and augment inflammatory or anti-inflammatory pathways, on early SIV infection of rhesus macaques. Our study revealed that similar to the GI mucosae, oral CD4+ T cells were rapidly depleted, and as one of the first comprehensive analyses of the oral microflora in SIV infection, we also observed significant modulation among two genera, *Porphyromonas* and *Actinobacillus,* early after infection. Interestingly, although Pbx therapy did not substantially protect against oral dysbiosis or ameliorate cell loss, it did dampen inflammation and T cell activation. Collectively, these data provide one of the most comprehensive evaluations of SIV-induced changes in oral microbiome and CD4+ T cell populations, and also suggest that oral Pbx could be a simple therapy to improve anti-inflammatory states in addition to more traditional antivirals.

## INTRODUCTION

Pathogenic HIV and SIV infections are characterized by early and sustained loss of mucosal homeostasis, inflammation, loss of CD4+ T cells and translocation of microbial products from the intestines into circulation and lymphoid tissues ^1–4^. Circulating lipopolysaccharide (LPS) and sCD14 are established biomarkers for microbial translocation and major modulators of inflammation by engaging TLRs. One proposed methodology to modulate inflammation is by regulating the microbiome of the lower gastrointestinal tract through probiotic supplementation (Pbx). Indeed, Pbx therapy has previously been used to improve general mucosal function and decrease the incidence of GI inflammation and disease, specifically inflammatory bowel disease, ulcerative colitis and Crohn’s disease ^5–11^. Pbx enhance NF-kB signaling and improve epithelial barrier function, likely through TLR signaling ^12–14^, and multiple lines of evidence suggest Pbx use in persons living with HIV (PLWH) may reduce microbial translocation and inflammation ^15–18^. Further studies using SIV infected nonhuman primate models also demonstrate improved immune function, reduced cellular immune activation, and decreased co-morbid conditions ^19,20^.

Due to a lack of global access to cART and ongoing oral immune activation in PLWH, oral complications in HIV disease are increasingly common. Oral bacterial and mycotic co-infections such as candidiasis, necrotizing gingivitis, and periodontitis are all common in PLWH ^21–23^, as are KSHV and EBV infection of the oral epithelia resulting in increased oral tumors and hairy leukoplakia ^24–26^. Much like has been shown in HIV disease, complications of the oral mucosa are also routinely found during experimental SIV infection of macaques. Although the mechanisms are not entirely clear for oral diseases in HIV/SIV infections, SIV-induced dysbiosis of the oral microbiome does result in increased production of cytokines and other soluble inflammatory mediators in the oral cavity, while simultaneously suppressing antimicrobial functions and epithelial development ^27^.

So called innate lymphoid cells (ILC) are identifiable by their lack of common lineage markers (those identifying T cells, B cells, NK cells, and myeloid cells), and can be distinguished by high expression of the IL-7 receptor, CD127 ^28–31^. ILC3 in particular play a major role in modulating gut homeostasis, but can be disrupted by changes in the inflammatory microenvironment, infectious disease, and shifts in the microbiome ^32^. HIV and SIV infections drive a perturbation of ILC3 resulting in loss of intestinal barrier integrity, increased inflammation and potential dysregulation of the local microflora. Specifically, others and we have shown that acute SIV infection causes depletion of ILC3 in the GI mucosae ^33–35^. In PLWH, ILCs are depleted but partially restored by ART ^36^. Much like had been shown in SIV-infected macaque models loss of ILC3 is associated with microbial translocation. Neither HIV nor SIV infection directly depletes ILC3, but rather changes in the cytokine milieu and local microbiotic signals lead to increased apoptosis ^33,34,36^. Importantly, we have previously shown that probiotic therapy improves IL-23 production and as a result expands ILC3 numbers in naïve macaques ^37^.

It has been shown that HIV alters human microbiome including oral bacteria communities ^38^. Periodontal disease, a common oral inflammatory infection in HIV infected patients, has been shown to be associated with general dysbiosis in the oral cavity, as well as with HIV disease progression ^39^. Dysbiosis in the oral cavity and changes to the oral bacterial communities might play a role in the sustained host proinflammatory responses to the virus. Restorative therapies that could improve oral and GI function, inflammation, and microbial health could be of significant benefit to PLWH. Indeed, previous studies by our groups have demonstrated that therapy with probiotic, beneficial bacteria can improve mucosal health and dampen inflammation in health macaques as well as in the context of SIV infection ^37,40,41^. Thus, we hypothesized that probiotic supplementation prior to SIV infection and during acute infection may enhance the oral microbiome and prevent pathogenic consequences of SIV. In this study we evaluated the effects of pre- and acute administration of oral probiotic therapy on viral kinetics, inflammation, CD4+ T cell loss, microbiome, and ILC3 modulation in the oral and GI mucosae, and associated lymphoid tissues in SIV-infected rhesus macaques.

## RESULTS and DISCUSSION

### Early probiotic administration during SIV infection

To determine the effects of probiotic (Pbx) administration pre-SIV and during acute SIV infection, we evaluated 12 Indian origin rhesus macaques, 6 receiving oral Pbx and 6 receiving oral vehicle control. Animals were given Pbx for a total of 42 days: 28 days pre-virus challenge and for a subsequent 14 days following challenge (**Figure 1A**). Macaques were intrarectally challenged with SIVmac251 as described previously ^42^, and buccal, colorectal, and lymph node biopsies in addition to blood draws were collected as indicated. No significant differences were observed in plasma viral loads longitudinally (**Figure 1B-C**), nor were any differences observed in peak viral loads between the two groups (**Figure 1D**). However, previous studies have also reported that the benefits of probiotic therapy in HIV an SIV are uncoupled from viremia which is generally not affected ^40^.

**Figure 1.**
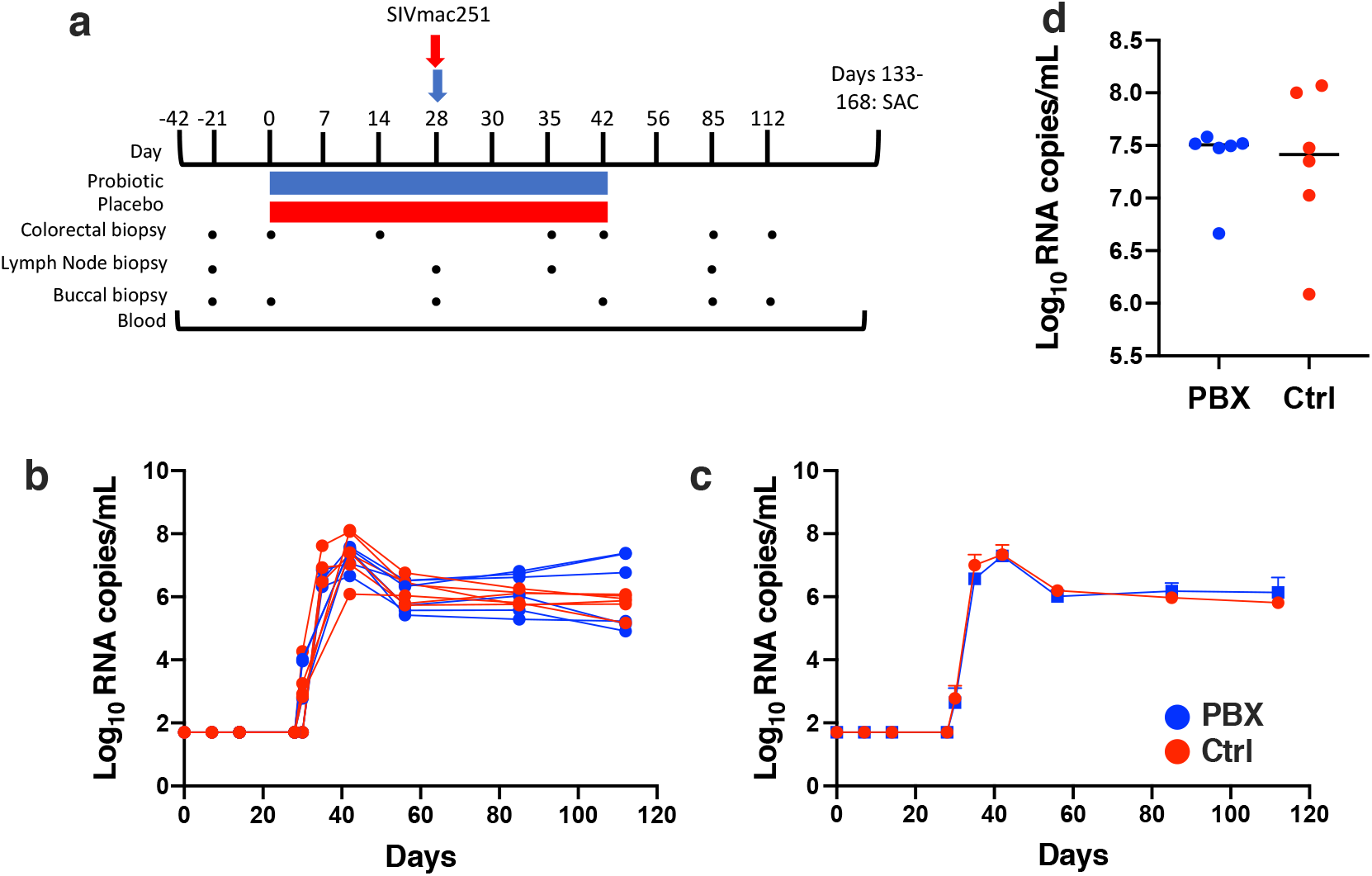
Oral Pbx therapy and SIVmac251 challenge in rhesus macaques. Timeline of study protocol (A); Pbx (blue) and vehicle control (red) groups. Both groups were challenged intrarectally with SIVmac251 at day 28. Tissue collections and treatment duration are indicated. Longitudinal viral loads (B) for individual animals, mean longitudinal (C) viral loads, and peak viral loads (D) of probiotic and control groups are quantified as described in the Methods.

### T cell responses during Pbx treatment of SIV-infected macaques

To assess broad antiviral T cell responses we performed longitudinal ELISPOT assays on PBMC as well as tissues at necropsy. Interestingly, Pbx-treated animals consistently had lower levels of both Gag (4-fold at day 112) and Env (9-fold at Day 112) ELISPOT responses (**Figure 2A-B**). Further, the total magnitude of the ELISPOT response (as measured Pol, Env, Gag) in control animals was significantly greater at necropsy in all tissues measured (PBMC, spleen, oral lymph nodes, and tonsil).

**Figure 2.**
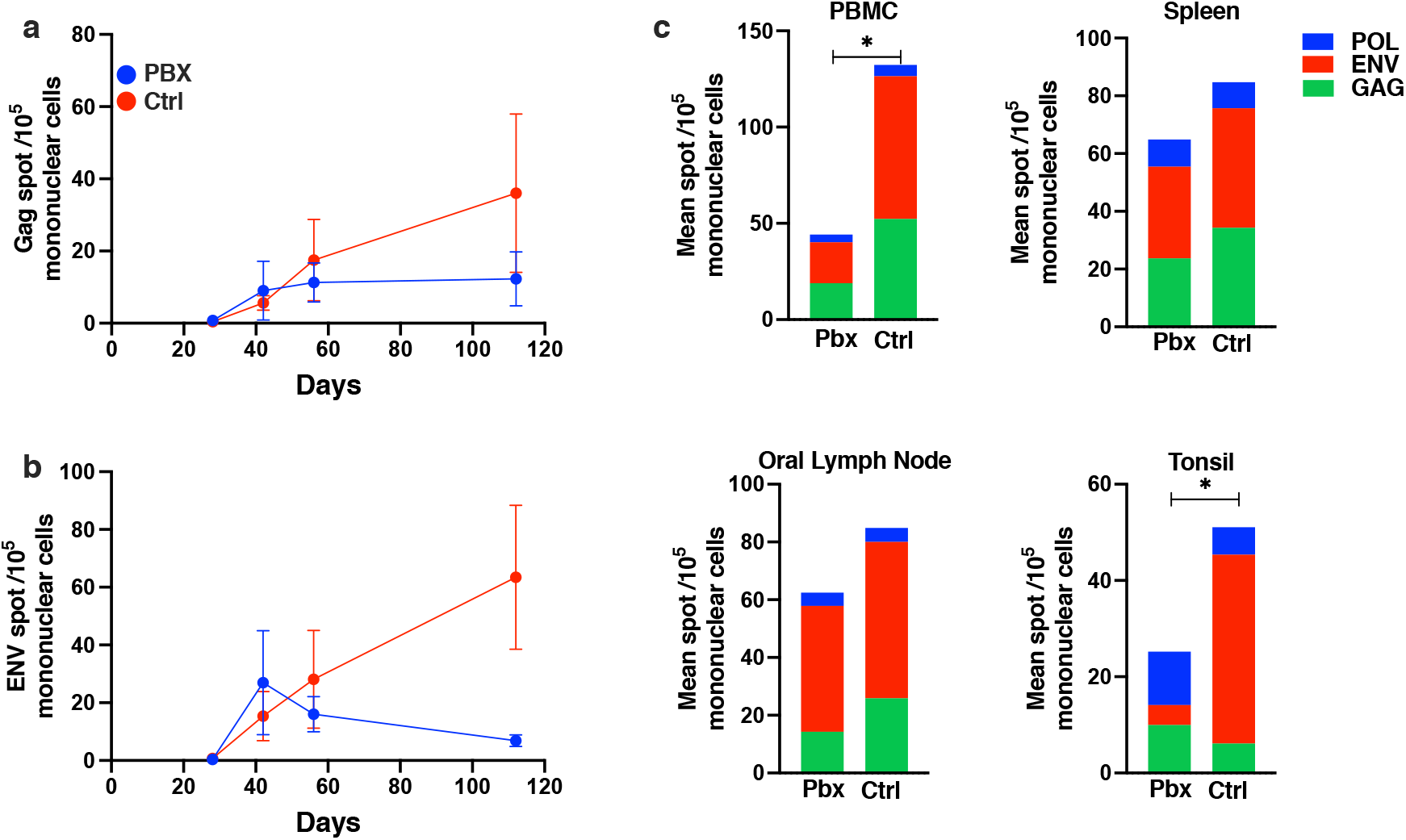
ELISPOT responses following SIV challenge. Mean spot forming cells using GAG (A) and ENV (B) peptide pools analyzed longitudinally following SIVmac251 infection. PBMC, spleen, oral lymph node, and tonsil mean spot forming cells (C) from probiotic and control animals are shown for POL (blue), ENV (red), GAG (green) ELISPOTS at necropsy. Unpaired *t* tests were used to compare total ELISPOT values between groups; *, P < 0.05.

Since SIV infection has a significant impact on memory CD4+ T cell populations, as well as CD8+ T cell activation, we next used polychromatic flow cytometry to evaluate these cell subsets in detail in Pbx-treated animals. As expected, the frequency of total CD4+ T cells in circulation declined early following SIV challenge concomitant with an increase in the frequency of CD8+ T cells (**Figure 3A**). Similar trends were observed in colorectal tissue where the shift was even more robust (**Figure 3B**), in line with previous studies ^43,44^. Remarkably, this study demonstrated for the first time an acute and sustained loss of CD4+ T cells in the buccal mucosa following SIV infection (**Figure 3C**). However, little to no difference was observed in CD4+ and CD8+ T cell dynamics in any tissue when comparing Pbx-treated to control animals. There was also little difference between the two groups when specifically examining CD28+CD95+ central memory CD4+ T cells in any tissue (**Figure 4A**). However, Pbx treatment reduced Ki67 expression (measured at day 42 Pbx/Day 14 post-challenge) in both bulk and CM CD4+ T cells in most tissues (**Figure 4B**). This suggests that while Pbx may not protect CD4+ T cell loss, it may reduce T cell activation, which is well in line with previous observations ^37^. These data indicate Pbx could be a beneficial augmentive therapy to protect CD4+ T cells and reduce overall cellular activation.

**Figure 3.**
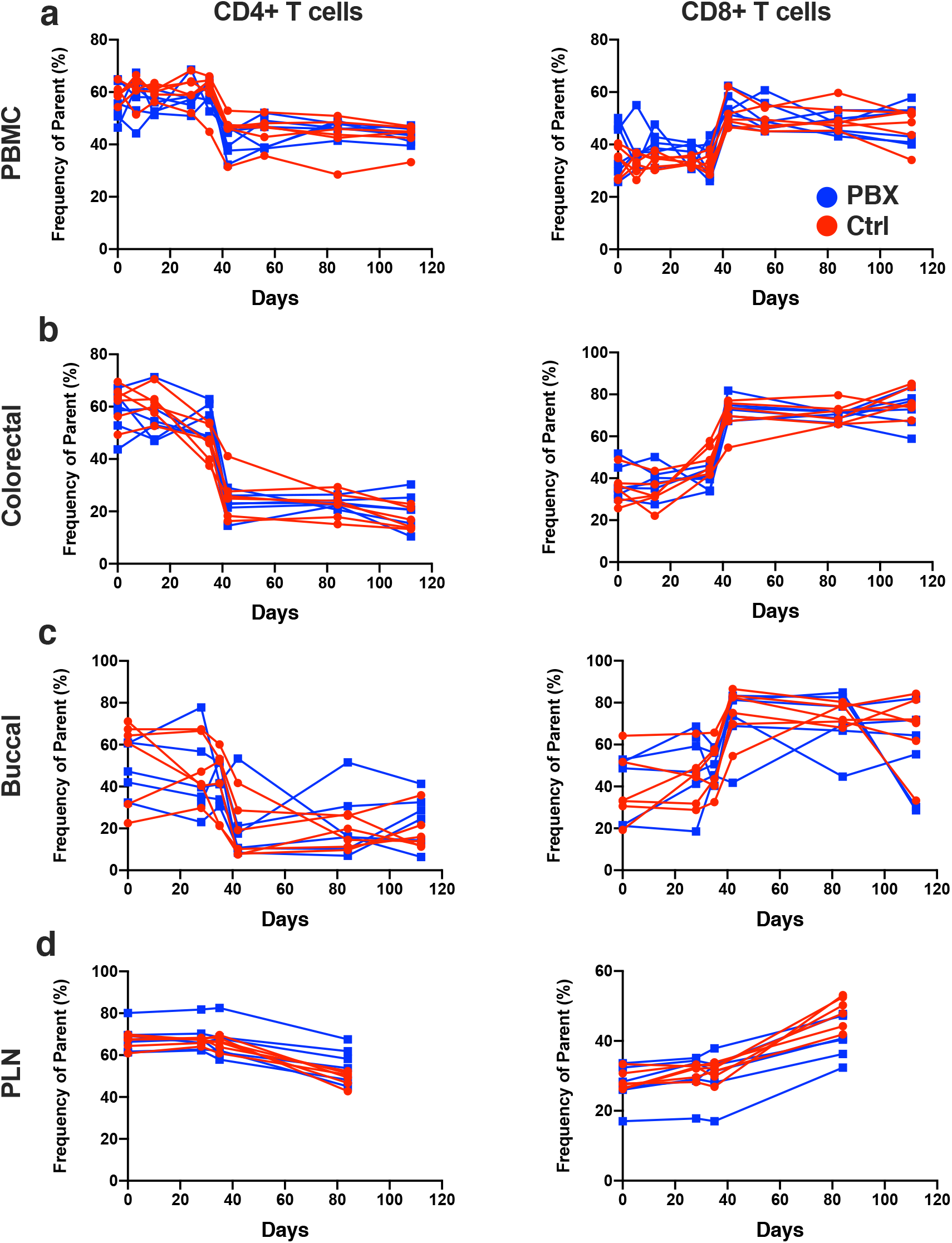
Bulk T cell analyses following Pbx therapy and SIV challenge. Longitudinal frequencies of CD4+ and CD8+ T cells in PBMC (A), colorectal (B), buccal (C), and PLN biopsies (D) are compared between control (red) and Pbx groups (blue). PLN, peripheral lymph nodes.

**Figure 4.**
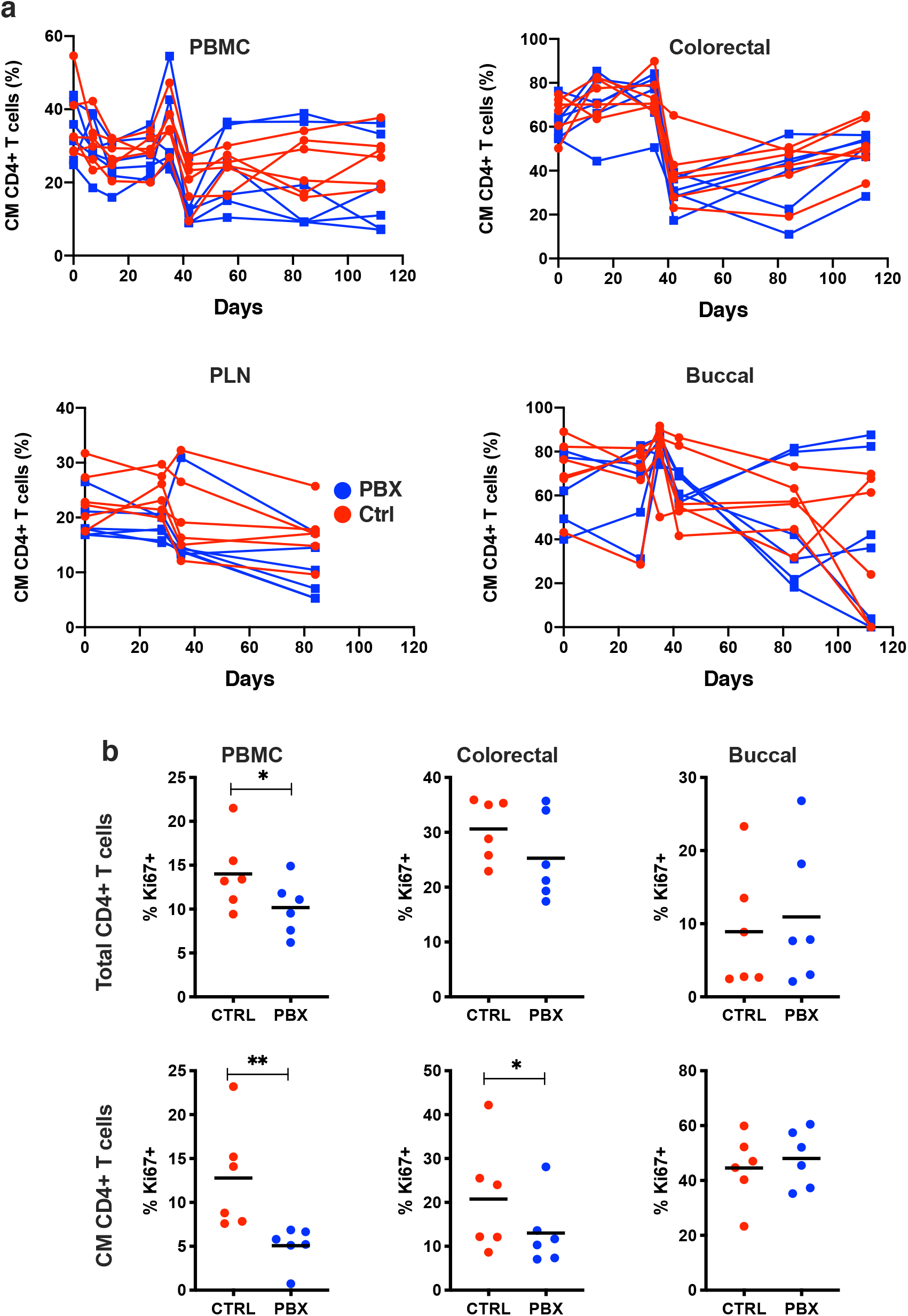
Memory CD4+ T cell analyses following Pbx therapy and SIV challenge. Central memory CD4+ T cells longitudinal frequency (A) for PBMC, colorectal, peripheral lymph node (PLN), and buccal, comparison of probiotic and control groups. Expression of Ki-67 on CD4+T cells and central memory T cells (B) between probiotic and control groups. Unpaired *t* tests were used to compare groups; *, P < 0.05; **, P < 0.01.

### Effects of probiotic treatment on ILC3 populations during SIV infection

We and others have previously shown that HIV and SIV infection can modulate ILC3 populations in the mucosae. Further, our groups have demonstrated that probiotic supplementation can actually expand ILC3 numbers in some animals and could possibly function as a putative rheostat mechanism of action for controlling inflammation during lentivirus infections. In this study we quantified the frequency of Lin-NKp44+NKG2A/C– ILC3, as described previously ^34,45^, in longitudinal colorectal and buccal biopsies. Interestingly, little change was observed in ILC3 frequencies during the probiotic supplementation period (day 0 to day 28) in this study (**Figure 5**). Although some animals demonstrated minor transient increases in ILC3 day 14 (colorectal) or day 28 (oral) following probiotic supplementation, these changes were not significant. It is unclear why ILC3 might not be expanded in this context, but previous work from our laboratory has shown that colorectal ILC3 are less likely to be increased compared to those in the jejunum and there is significant animal-to-animal variation ^37^. Further, Pbx only appeared to marginally impact SIV-induced changes in ILC3 frequencies, indicating that while Pbx may impact T cell responses, it is unlikely to protect against loss of ILC3.

**Figure 5.**
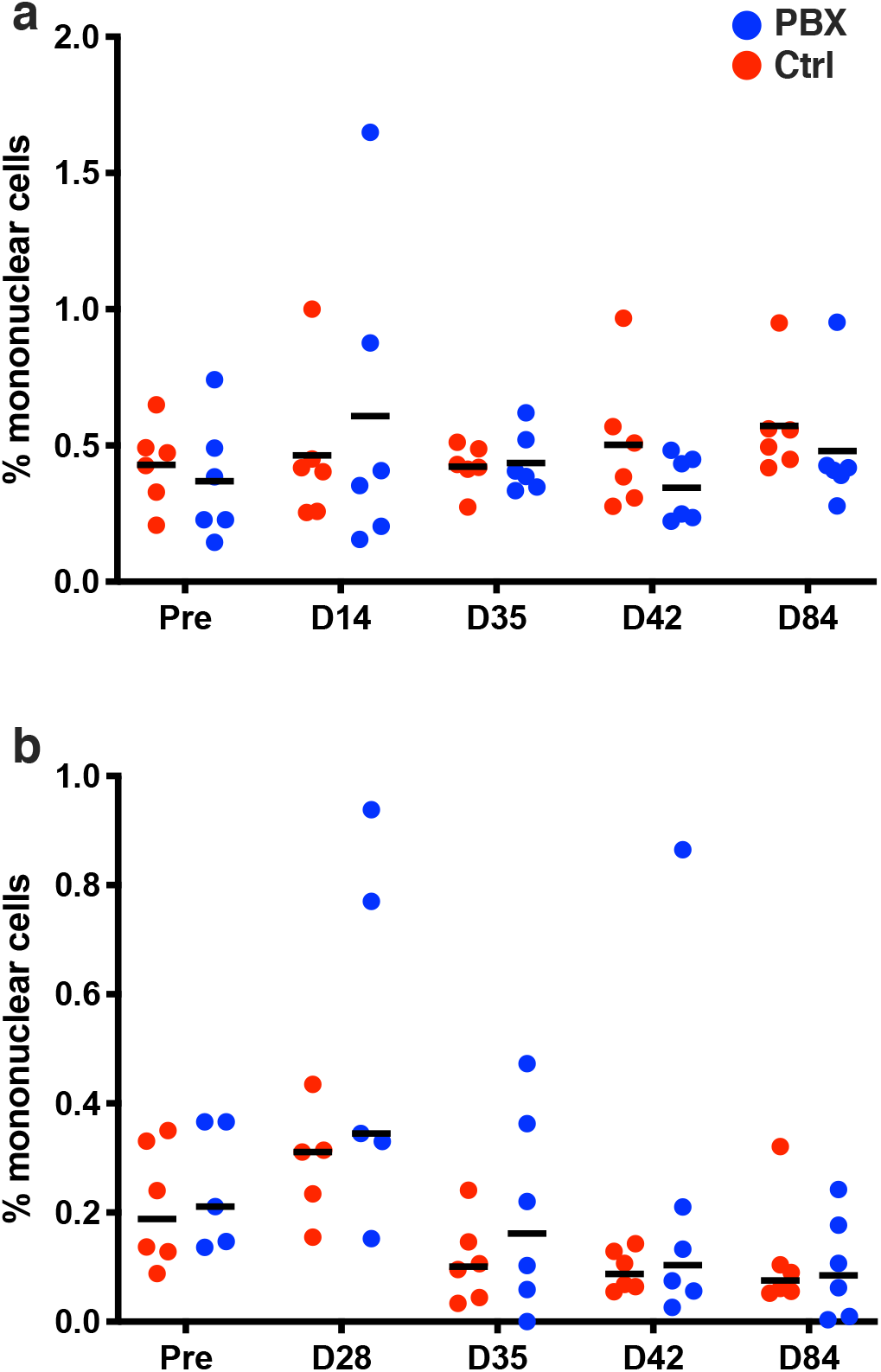
ILC3 analyses following Pbx therapy and SIV challenge. ILC3 frequencies were quantified in colorectal (a) and buccal (b) biopsies are multiple timepoints (relative to PBX initiation) and during SIV infection (d28 challenge).

### Soluble measures of inflammation and immune activation

To further address the impact that Pbx and ILC3 modulation might be having on global inflammation during SIV infection we compared soluble inflammatory mediators in the circulation by Luminex. Animals treated with Pbx had a distinctly lower inflammatory profile on day 28 (day of challenge), suggesting that Pbx treatment alone was sufficient to decrease soluble mediators of inflammation (**Figure 6A**). This reduced inflammatory profile persisted from day 42 (day 14 post-challenge), and even persisted in some animals up to day 112 (day 84 post-challenge. However, Pbx therapy did not appear to have any effect on microbial translocation products (**Figure 6B**), which is similar to HIV studies showing improved immune function but no consistent decline in circulating bacterial products ^46,47^.

**Figure 6.**
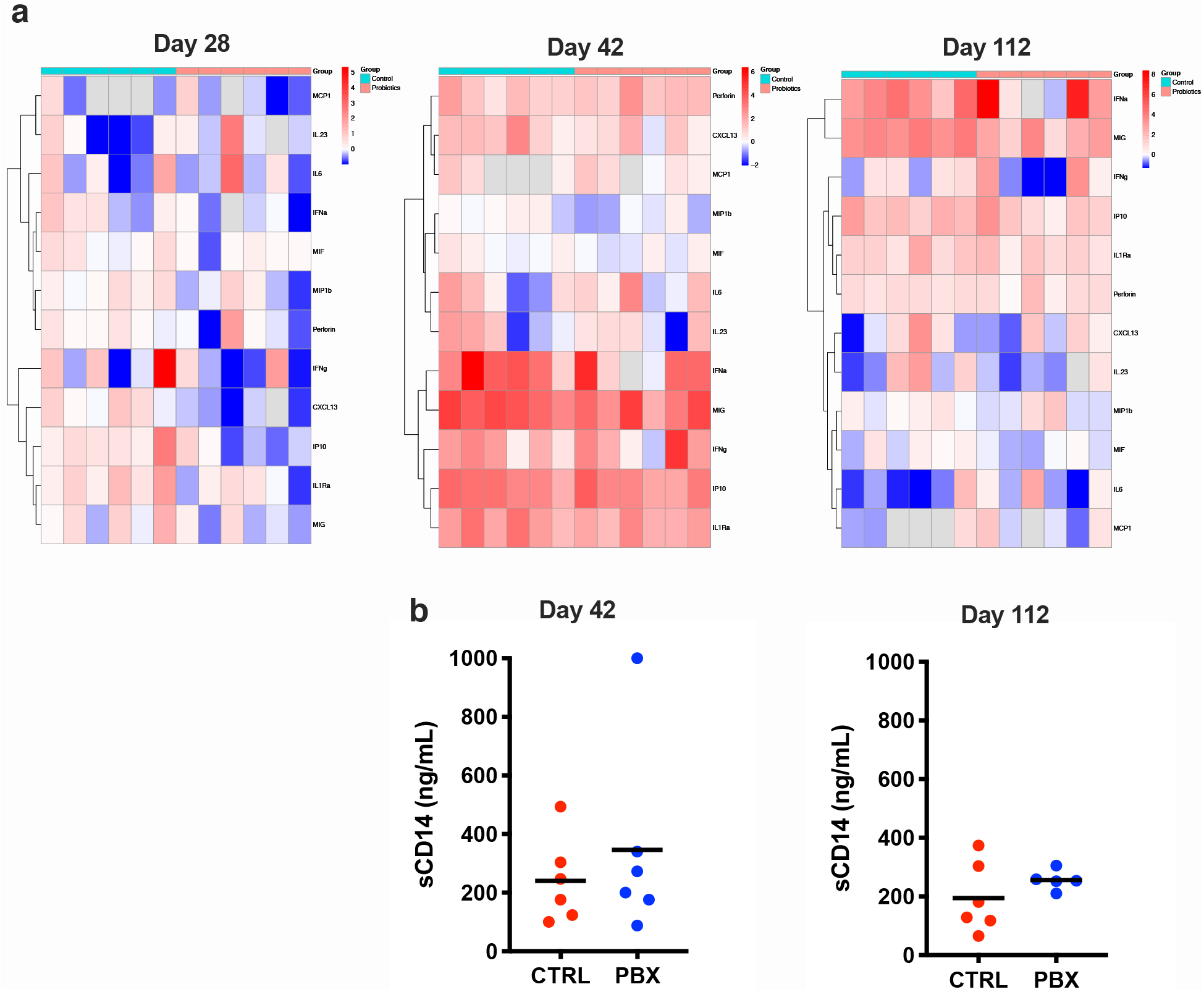
Soluble measures of inflammation. (A) Heatmaps of Luminex data of inflammatory cytokines present in both control and probiotic groups at days 28, 42, and 112 shown as log2 fold changes over baseline values. Columns correspond to individual macaques included in this study. (B) Microbial translocation products post infection of individual animals receiving placebos (red) and probiotics (blue) (B) taken at day 42 and 112.

### Probiotic supplementation partially modulates oral microbiome during SIV infection

To study the effects of Pbx on the microbiome, we used 16s rRNA gene sequencing to profile bacterial communities in the buccal and colonic mucosa. No notable differences in community richness, evenness, Shannon diversity or over all principal components analysis differences were observed between groups in either oral or gut samples, consistent with previous studies of probiotic supplementation. The most abundant bacteria in the oral mucosa of all animals were of Firmicutes phyla, with lesser representation of Bacteroidetes, Proteobacteria, Fusobacteria, and Actinobacteria (not shown). The colorectal mucosa top phlya was Epsilonbacteraeota followed by Bacteroidetes and Spirochaetes (not shown). At the phyla level, no notable differences between groups or across time points were observed in either site. At the genus level for the oral mucosa, a high abundance of *Streptococcus* and *Veillonella* was observed in all samples, followed by minor abundances of *Alloprevotella, Carnobacterium, Actinobacillus, Porphyromonas, Haemophilus, Gemella,* and *Prevotella* (**Figure 7A**). The top genera for colorectal samples were *dominated* by *Helicobacter* with an abundance of fifty-five percent followed by *Prevotella, Treponema, Alloprevotella, Ruminococcaceae, Camplobacter* with abundances starting at nine percent (**Figure 7B**). We performed differential abundance analysis on the Pbx and placebo groups across all timepoints in the oral mucosae. In the placebo group, *Porphyromonas* and *Actinobacillus* were found to be significant between day −21 and 0 (*p*=0.012 and 0.021), day −21 and day 28 (*p*=0.028), and day 28 and day 42 (*p*=0.004 and 0.001) (**Figure 7C-D**). In the Pbx group, *Porphyromonas* and *Actinobacillus* were significant between day −21 and day 28 (*p*=0.04) and day 42 and day 112 (*p*=0.037 and *p*=0.017) (**Figure 7D**). However, notable differences were not observed between groups, however it does appear that the changes observed were dampened in the probiotics group compared to the placebo group. Thus, while probiotics did not have a substantial impact in modulation of the microbiome, these changes reflect possible dampening of SIV-mediated dysbiosis in the oral microbiome.

**Figure 7.**
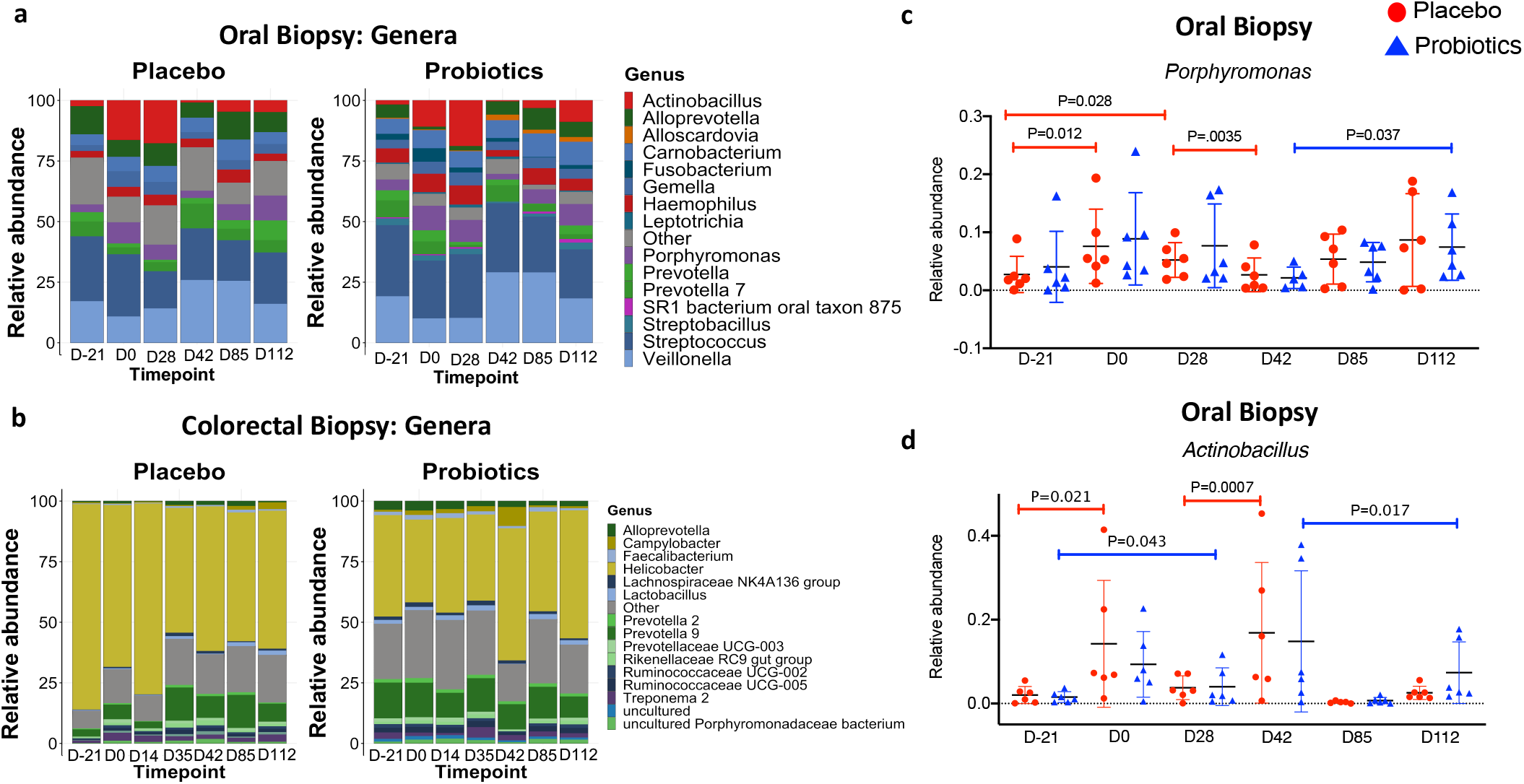
Microbiome alterations following Pbx therapy and SIV challenge. (A) Taxonomic barplot of average relative abundance for top 15 genera level bacteria separated by experimental group across timepoints for buccal biopsies. (B) Taxonomic barplot of average relative abundance for top 15 genera level bacteria separated by experimental group across timepoints for colorectal biopsies. (C) Scatter plot of relative abundance of *Porphyromonas for buccal biopsies* separated by experimental group (red is placebo and blue are probiotics) across all timepoints. DESeq2 was used for statistical assessment and adjusted p values of <0.05 significant. (D) Scatter plot of relative abundance of Actinobacillus for buccal biopsies separated by experimental group (red is placebo and blue are probiotics) across all timepoints across timepoints. DESeq2 was used for statistical assessment and adjusted p values of <0.05 significant.

### Concluding remarks

In summary our study presents a number of new findings relevant to SIV immunobiology. First, SIV infection clearly demonstrates an early depletion of CD4+ memory T cells in the oral mucosa. Next, Pbx does not appear to protect against loss of CD4+ T cells or ILC3 but does reduce measures of systemic inflammation. These data also offer one of the most in-depth characterizations of microbiome in the oral mucosae, where we demonstrated novel changes to the oral microbiome after SIV infection, which are slightly dampened with probiotics. However, no substantial microbiome alterations were observed between probiotics and placebo groups. Overall, our studies indicate a role for oral Pbx in modulating SIV-induced inflammation, but additional work will be required to tease out detailed mechanisms of action.

## METHODS

### Ethics statement

All animals were housed at Biomere, Inc., Worcester, MA in accordance with the rules and regulations of the Committee on the Care and Use of Laboratory Animal Resources. Animals were kept in agreement with the American Association for Accreditation of Laboratory Animal Care standards and *The Guide for the Care and Use of Laboratory Animals*^39^. Animals were fed standard monkey chow diet supplemented daily with fruit and vegetables and water ad libitum. Social enrichment was delivered and overseen by veterinary staff and overall animal health was monitored daily. Animals showing significant signs of weight loss, disease or distress were evaluated clinically and then provided dietary supplementation, analgesics and/or therapeutics as necessary. Animals were humanely euthanized using an overdose of barbiturates according to the guidelines of the American Veterinary Medical Association. All studies reported here were performed under IACUC protocols 17-02 and 16-08 which was reviewed and approved by the Biomere IACUC. When necessary, macaques were immobilized with ketamine HCl (Parke-Davis) at approximately 10 mg/kg and injected intramuscularly after overnight fasting. Blood samples were collected using venipuncture.

### Animals, probiotic supplementation and SIV challenge

Sixteen experimentally naive age-matched Indian-origin rhesus macaques (*Macaca mulatta*) were analyzed in this study. All animals were colony-housed at Biomere, Inc. and were free of simian retrovirus type D and simian T-lymphotropic virus type 1. Animals were challenged intrarectally with SIVmac251 as described previously ^42^. Six animals received Visbiome for 42 days mixed with food, while six received food only (vehicle) control. No adverse events due to Pbx therapy were observed in any of the animals.

### Blood and tissue processing

Blood samples were collected in EDTA-treated tubes, and peripheral mononuclear cells (PBMCs) were isolated using standard density gradient centrifugation over lymphocyte separation media (MP Biomedicals, Solon, OH) and any contaminating red blood cells were lysed using hypotonic ammonium chloride solution. Processing of tissues was carried out using protocols optimized in our laboratory^27,30,41^. Briefly, colon and buccal tissue were cut into 1 cm^2^ then incubated in 5mM EDTA for 30 min before undergoing mechanical and enzymatic disruption. Samples were then gently pushed through filters before lymphocytes were isolated via bilayer (35%/60%) isotonic Percoll density gradient. Lymph nodes and spleen were trimmed of excess tissue then mechanically disrupted. Manual cell counts were performed for each sample using Trypan Blue. Two million cells from each sample were then used for real-time flow cytometry staining while remaining cells were cryopreserved in a DMSO solution and stored in liquid nitrogen vapor.

### ELISPOT

ELISPOTs were performed with Mabtech IFN-y ELISpot Plus Kits. Peptides used were SIVmac239 Gag (Cat: 12364), ENV (Cat: 12635), and POL (Cat: 12766) obtained through the NIH AIDS Reagent Repository and aliquoted into pools. Briefly, samples were thawed and rested for four hours. Cells were then counted and aliquoted into 96 well plates at 0.25M-0.2M cells per well. Following incubation at 37 degree C and substrate development, plates were read by ZellNet Consulting (Fort Lee, NJ).

### Antibodies and flow cytometry

Flow cytometry staining of mononuclear cells was carried out for cell surface and intracellular molecules (**Supplementary Table 1**). Briefly, samples were incubated with the LIVE/DEAD Aqua amine dye (Invitrogen, Carlsbad, CA), then washed before staining with surface antibodies. After staining, cells were permeabilized using Thermo Fisher Scientific Fix & Perm buffer kit (ThermoFisher Scientific, Waltham, MA) and incubated with the intracellular staining antibodies. Isotype-matched controls were included in all assays. All acquisitions were made on an LSRII (BD Biosciences) and analyzed using FlowJo (9.9.6).

### Viral load quantification

Quantified RNA was used for RT-PCR against a conserved region of *gag* using gene-specific primers ^48^. In the first step, RNA was reverse-transcribed followed by treatment with RNase for 20 min at 37°C. Next, cDNA was amplified using 7300 ABI Real-Time PCR system (applied Biosystems) according to the manufacturer’s protocol.

### DNA extraction, 16S rRNA gene sequencing and data analysis

We extracted DNA from cryopreserved buccal and colorectal biopsies using the PowerFecal DNA Isolation Kit (Qiagen, Valencia, CA). We then prepared sequencing libraries as described by the Earth Microbiome Project and sequenced them using the Illumina MiSeq Sequencer (Illumina, San Diego, CA). 515F-806R primers were used to sequence the V3-V4 region of the 16s SSU rRNA as previously described ^49^. 16s sequencing reads were demultiplexed, trimmed and then processed using QIIME2 (version 2020.2) ^50^. Operational taxonomic units (OTUs) were clustered at 99% similarity using the DADA2 method and assigned taxonomy with the Silva 132 classifier for taxonomic determination. All sequence reads and operational taxonomic unit (OTU) observations were included in our analyses, in order to maximize the observed diversity of the bacterial communities and was done utilizing R (v3.6.0) packages. Alpha diversity (richness, evenness (Pielou)), and Shannon diversity) was calculated by the microbiome package, beta diversity was calculated with pairwise sample dissimilarity and unweighted unifrac ordination analysis by means of principal coordinates analysis (PCoA) using the vegan package for Ordination, Diversity and Dissimilarities. Statistical analyses were completed using the Pairwise Adonis package. Differential abundance analysis was completed using DESeq2. Beta diversity and taxonomic plots were created utilizing the phyloseq and ggplot. Alpha diversity and differential abundance plots were created in Prism 8 version 8.4.3 (GraphPad Software).

### Statistical analyses

Statistical analyses were carried out using Prism Version 8.3.0 (GraphPad Software). Unpaired, nonparametric, Mann-Whitney *U* tests, Wilcoxon, and *t* tests were used where indicated, and P<0.05 were assumed to be significant. Samples were grouped and heatmaps were generated in Prism version 7.0d (GraphPad Software). For beta diversity, statistical significance was assessed using the pairwise permutational multivariate analysis of variance (PERMANOVA; Adonis) test in R. For taxonomic differential abundance DESeq2 was used and accounted for sequencing depth and multiple comparisons using the Benjamini and Hochberg method to control the false discovery rate (FDR) and adjusted p values of <0.05 were considered significant.

### Multiplex cytokine analysis of plasma

Plasma samples were analyzed for rhesus macaque cytokines and chemokines by Luminex using established protocols for non-human primates. Evaluation of analytes IL-6, IL-12p70, IL-17, TNF-α, IL-2, G-CSF, CXCL13, MIP-1β, IP-10, IL-1Ra, Perforin, IFN-α, IFN-γ, MCP-1, MIG, Il-23 and MIF were included in this assay. Only analytes quantifiable above the limit of detection in plasma are presented. Log2 fold change for Luminex data was calculated per RM in this study. Briefly, each rhesus macaque timepoint log2 fold change was calculated using the following formula:

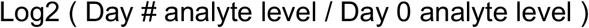

NaN or Inf values were replaced with NA. Log2 fold change was calculated in R v4.0.0 (https://www.R-project.org) and heatmaps were generated with pheatmap v. 1.0.12 (https://CRAN.R-project.org/package=pheatmap). Row clustering was performed on cytokine log2 fold changes with default parameters. Column clustering was not performed.

## Acknowledgements

The authors thank Michelle Lifton, Dr. James Whitney, and Dr. So-yon Lim for technical assistance, Dr. Dan Barouch for providing the SIV challenge stock, as well as Dr. Luis Giavedoni (Southwest National Primate Research Center) for assistance with Luminex assays. We also thank Dr. Angela Carville and Biomere, Inc. for animal management.

## Author contributions

R.K.R and N.K. designed the project. R.J., S.S., K.K., V.V., B.H., S.V.S., D.R., and C.M processed samples. V.V. coordinated animal studies, and R.J., K.K., C.B., L.S., and R.K.R. analyzed the data. All authors contributed to the final preparation of the manuscript.

---

**Supplementary Figure 1.**
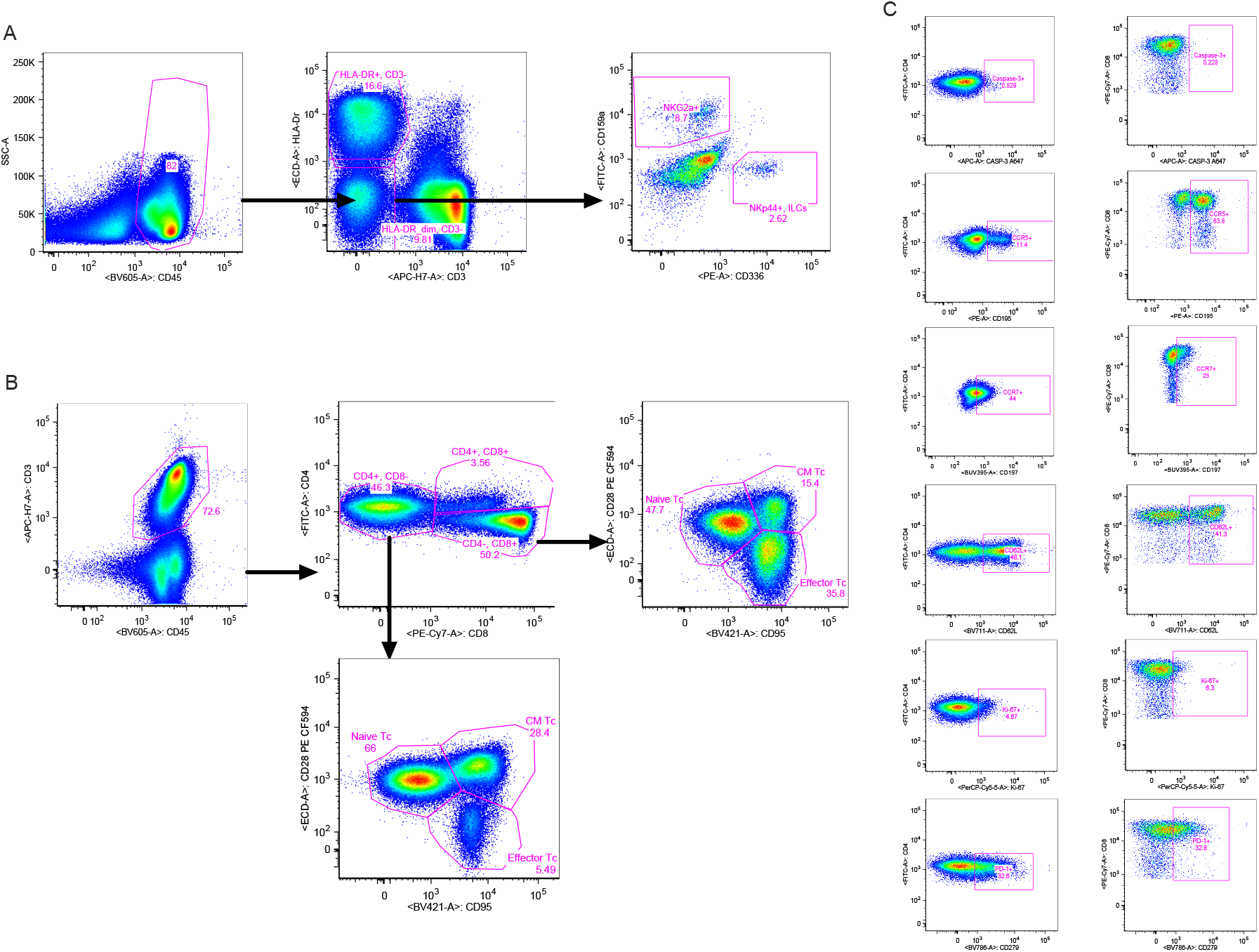
Representative flow cytometry gating. Representative gating strategies are shown for (A) NKG2A/C+ NK cells and NKp44+ ILC3, (B) T cells, and (C) T cell phenotypes.

**Supplementary Table 1.**

| Target | Fluorochrome | Clone | Vendor |
| --- | --- | --- | --- |
| Caspase-3 | Alexa647 | C92-605 | BD Pharmingen |
| CD3 | APC CY7 | SP34.2 | BD Pharmingen |
| CD4 | FITC | L200 | BD Pharmingen |
| CD8 | PAC BLUE | RPA-T8 | BD Pharmingen |
| CD8 | PE CY7 | RPA-T8 | BD Pharmingen |
| CD11C | A700 | 3.9 | Affymetrix |
| CD14 | BV650 | M5E2 | BD Pharmingen |
| CD16 | BUV496 | 3G8 | BD Pharmingen |
| CD20 | BUV395 | L27 | BD Pharmingen |
| CD28 | PE CF594 | 28.2 | BD Pharmingen |
| CD45 | BV605 | D058-1283 | BD Pharmingen |
| CD56 | BV786 | NCAM16.2 | BD Pharmingen |
| CD62L | BV711 | SK11 | BD Pharmingen |
| CD95 | BV421 | DX2 | BD Pharmingen |
| CD123 | PE CY7 | 7G3 | BD Pharmingen |
| CD159A | FITC | REA110 | MILTENYI |
| CD195 | PE | 3A9 | BD Pharmingen |
| CD197 | BUV395 | 150503 | BD Pharmingen |
| CD279 | BV786 | EH12.1 | BD Pharmingen |
| CD336 | PE | Z231 | BECKMAN COULTER |
| HLA-DR | ECD | IMMU-357 | BECKMAN COULTER |
| Ki-67 | PERCP CY5.5 | B56 | BD Pharmingen |

